# Rprot-Vec: A deep learning approach for fast protein structure similarity calculation

**DOI:** 10.1101/2025.01.25.634852

**Authors:** Yichuan Zhang, Wen Zhang

## Abstract

The prediction of structural similarity and homology detection of protein sequences has always been a very challenging and valuable research. Based on protein structure similarity, rapid and accurate detection of homologous proteins can infer the functions of newly discovered or unannotated proteins. If we can use a method to quickly predict the structural similarity between two proteins, we can achieve the task of homologous detection of protein sequences. By using this method with partially known protein structures, we can quickly infer the functions of more unknown proteins. In this article, we propose an enhanced model Rprot-vec (Rapid Protein Vector) based on deep learning, which improves existing methods. It can quickly calculate the structural similarity of proteins and perform protein homology detection using only protein sequences. Meanwhile, our model can also be used to simulate proteins to accomplish more downstream tasks related to proteins. Our model performs well within the very important TM-score range, with an accurate similarity prediction rate of 65.3% for homologous proteins. In addition, for all protein pairs within the TM-score range, the average prediction error is 0.0561, indicating good performance. The superiority of our model can be demonstrated from various aspects. In addition, we have also supplemented the dataset in this field, providing data support for future researchers studying this field simultaneously.

**Supplementary information:** https://github.com/SuperZyccc/RProt-vec/

## 1 Introduction

Protein structure similarity prediction is an important direction in protein-related research, with the core objective of evaluating the three-dimensional structural similarity of two proteins through computational methods, enabling more downstream analysis topics.

By constructing a protein structure similarity prediction model, we can model proteins and represent different proteins using multidimensional vectors. At the same time, we can also study the evolutionary relationship of proteins. Even if the sequences of two proteins are significantly different, if their structures are similar, these two proteins have similar functions and may have a common evolutionary origin (*Zhang and Skolnick, 2005*).

In addition, we can also predict unknown protein functions through this study. Currently, many studies have shown that the structure and function of proteins are consistent (*Baker and Sali, 2001*) (*Orengo et al., 1999*). If a protein with an unknown function has a high structural similarity to a protein with a known function, then the protein with an unknown function is highly likely to possess that known function. We can also use this method to build a fast query protein coding database, which allows us to quickly screen and analyze structurally similar related proteins from massive data, making subsequent queries and research more efficient.

The research on efficient and automated tools for comparing the three-dimensional structures of proteins can be traced back to the 1990s, when the direct analysis was conducted by comparing the three-dimensional structures of proteins. Holm et al. proposed the Dali method in 1995, which is a dynamic programming algorithm that calculates the distance relationship between main chain atoms in the three-dimensional structure of proteins to find the optimal alignment (*Holm and Sander, 1995*). The DALI method uses Z-score as a similarity indicator to evaluate the significance of the comparison results. A high Z-score indicates that the comparison results are statistically significant, indicating that the structures of two proteins have a high degree of similarity. The core algorithm idea of DALI is still borrowed by many subsequent algorithms, laying the foundation for the modern algorithm framework of protein structure alignment.

Shindyalov et al. proposed the CE (Combinatorial Extension) method in 1998, which combines and extends local structural fragments of proteins to construct global structural alignment (*Shindyalov and Bourne, 1998*). Compared to the DALI method, which focuses more on global alignment, the CE method achieves a balance between structural flexibility and accuracy by gradually aligning globally through local alignment. The CE method uses RMSD as its similarity evaluation metric. RMSD represents the average deviation of atomic pairs after aligning two proteins. A smaller RMSD value indicates higher structural similarity. The fragment extension approach of the CE method also provides a way of thinking for subsequent related research.

Zhang et al. proposed the TM-align method in 2005, which can quickly and accurately compare the three-dimensional structural similarity of two proteins (*Zhang and Skolnick, 2005*). The TM-align method can capture both global alignment and local structural features of protein structures simultaneously, and uses TM-score as an evaluation metric to enable fair comparison of proteins of different lengths. Therefore, the TM-align method has been widely applied in protein structural biology research, including protein function prediction, protein evolutionary relationship analysis, and other fields.

In recent years, with the continuous development of artificial intelligence technology, it has gradually been applied to various fields of scientific research. As an important technology in the field of artificial intelligence, deep learning is currently widely used. The core of deep learning is to process complex data and tasks by simulating the neural network of the human brain. It has a multi-layer structure of artificial neural networks, which use stacking to extract features from massive training data and gradually learn the rules of the training data. Therefore, deep learning is a method with powerful representation and automated feature extraction capabilities.

In the prediction of protein structure similarity, traditional methods used to rely on protein structure data for calculation. However, according to the protein database PDB, the proportion of proteins with known three-dimensional structures in the total number of proteins is less than 0.1%, and the vast majority of protein sequences have known gene codes (*Burley et al., 2017*). Therefore, using a model that can quickly predict protein structure similarity without the need to predict protein structure would be a good choice.

Hamassy, T. et al. proposed the TM-vec method in 2023, which predicts structural similarity by calculating the structural similarity score TM-score of two protein sequences (*Hamamsy et al., 2024*). This model uses the ProtT5 model for precoding, and the model structure adopts the encoder part of the transformer and the fully connected layer as the main body. The protein sequence will be encoded into a fixed-length vector through this model, and the cosine value of the protein pair will be calculated to obtain the final TM-score. TM-vec provides an efficient and accurate tool for predicting protein structure similarity and can perform prediction work without obtaining the protein structure, laying the foundation for research in this field.

In this paper, Rprot-vec model with faster training speed and suitable for smaller datasets is proposed. This method continues the idea of using protein sequence pairs directly to predict structural similarity in TM-vec and proposes the idea of using bidirectional GRU instead of Transformer blocks in smaller models, while using convolutional neural networks (CNN) to extract features from different dimensions. Due to the use of bidirectional GRU instead of Transformer blocks and the reduction of training dataset requirements, better training results can be achieved with smaller datasets, reducing training time. In terms of model performance, Rprot-vec has good performance on homologous protein pairs and can accurately identify and judge homologous proteins. In addition, the dataset is the most important part of training, and due to the fact that TM-vec has not publicly released the training dataset, its contribution of the dataset to subsequent related research and training work is relatively small. Therefore, this paper will also publicly disclose the dataset used for training, to facilitate the development of related research in the future.

### 2 Methods

The Rprot-vec model proposed in this paper uses ProtT5 (*Elnaggar et al., 2021*) as the encoder for protein sequences, encoding each amino acid according to context. The main architecture of the model consists of Bi-GRU (Bidirectional GRU) and multi-scale CNN. The Bi-GRU is responsible for further extracting contextual features of the entire sequence, while the multi-scale CNN uses convolution kernels of different sizes to attempt to extract local features of different sizes. The structure of the model is shown in Fig. 1.

**Fig. 1.**
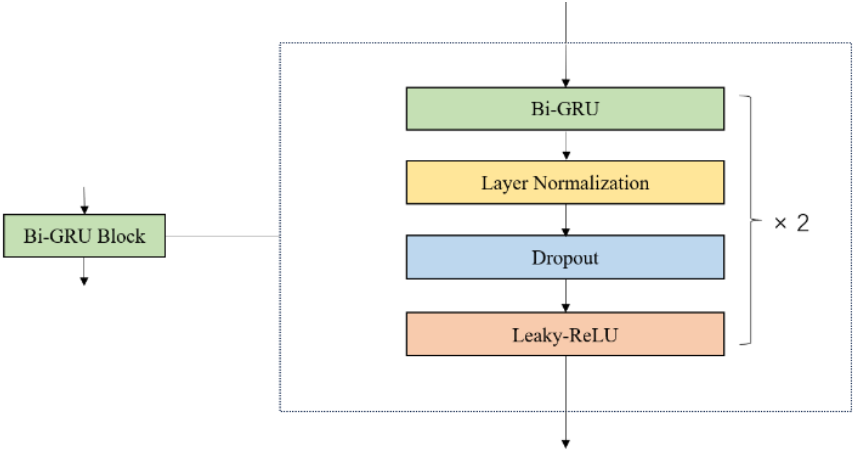
The model structure of the Rprot-vec

In our method, the ProtT5 model will receive the raw protein sequence and output a vector that converts each amino acid into a fixed dimension. Although there are only 20 standard types of amino acids, due to the influence of context, even the same two amino acids can be encoded into different vectors. By using ProtT5 as an upstream model, protein context features were extracted while encoding, providing rich feature inputs for downstream tasks.

In addition, we used Bi-GRU to extract global feature. Bi-GRU is an extension of standard GRU (*Cho, 2014*). Its main feature is that it can process sequences twice, once in the forward direction and once in the reverse direction. The advantage of doing so is that it can consider both the order from front to back and from back to front, which can better understand the bidirectional dependencies in sequential data. The internal structure diagram of the Bi-GRU block we use is shown in Fig. 2.

**Fig. 2.**
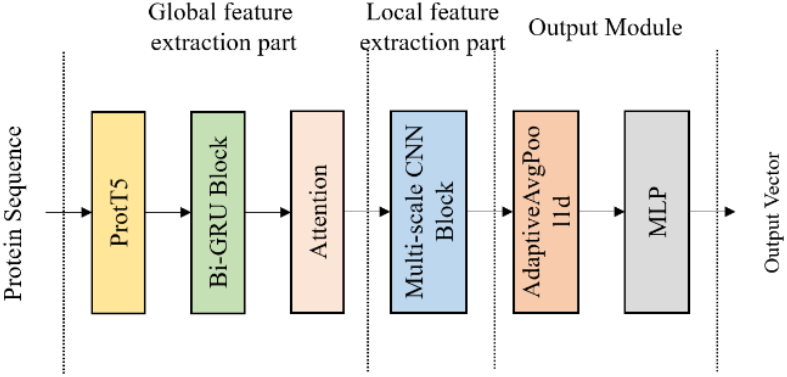
The internal structure of the Bi-GRU Block

After the Bi-GRU block, an attention layer is added. Through weighted aggregation, the amino acid features of each position can be directly associated with the amino acids of all other positions in the sequence, which compensates for the shortcomings of GRU and enhances the protein sequence processing capability of our model.

In protein sequences, the length of locally important amino acid fragments may vary, and we need to use a method to capture locally important features. Therefore, we adopted a multi-scale convolutional neural network, which can more effectively help us capture local features of protein sequences. The specific internal structure diagram of the multi-scale convolution block is shown in Figure 3. We used two different sizes of convolution kernels, 3 and 7, to help us capture local features of different sizes.

**Fig. 3.**
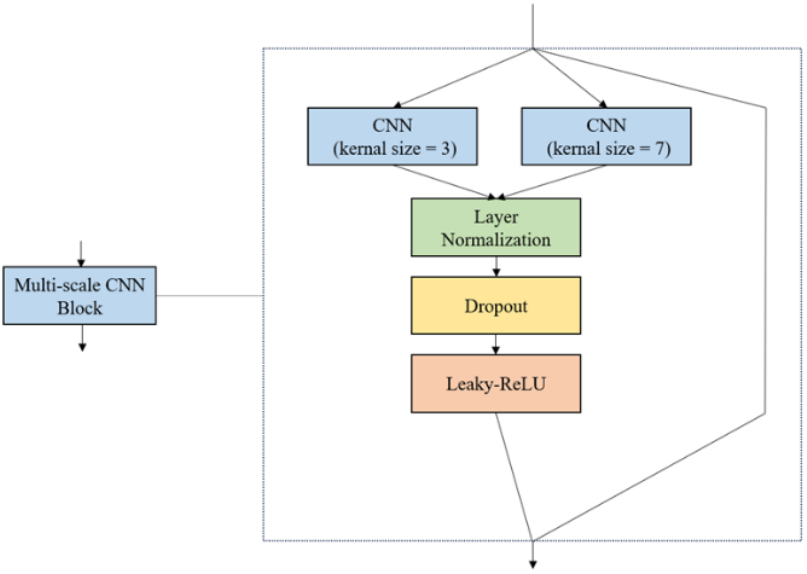
The internal structure of the Multi-scale CNN Block

We need to map the output of the convolutional layer onto a fixed dimensional vector, so we chose adaptive average pooling, which adjusts the length of each channel in the input sequence to a fixed target length and performs average pooling on the sequence during the process. Next, after passing through a fully connected layer, we obtain the fixed length vector *Output* that we need.

In the final similarity calculation section, we want to express the protein sequence in space using the output vector and also want to use this output vector to calculate the final protein structure similarity score TM-score. The positive half of cosine similarity ranges from 0 to 1. When the value is 1, it represents that the two spatial vectors completely overlap. When the value is 0, it represents that the two vectors are orthogonal. Therefore, using the positive half of cosine similarity as the final calculation metric is very suitable for our current situation. If two proteins with very similar structures are compared, we hope that their TM-score values will be close to 1, and we also hope that they will almost overlap in spatial vectors. And for proteins with completely dissimilar structures and no homologous information, we hope that their TM-score value is close to 0 and should appear orthogonal in spatial vectors.

## 3 Experiments

### 3.1 Dataset

In our paper, the dataset used for training was jointly generated based on the US-align method (*Zhang et al., 2022*) and the CATH protein database. Each dataset consists of three columns, namely protein sequence 1, protein sequence 2, and their protein structure similarity score TM-score.

#### (a) CATH database

The CATH database is a database dedicated to protein related research, named after four main levels: Class, Architecture, Topology, and Homologous Superfamily, which can be used to study the structure, function, and evolutionary relationships of proteins. The CATH database combines automated algorithms and manual annotation to ensure its accuracy and reliability (*Orengo et al., 1997*) (*Sillitoe et al., 2015*) (*Dawson et al., 2017*).

In our method, we first downloaded partial sequence data of known protein three-dimensional structures. Through automated script code, we automatically downloaded the corresponding sequence three-dimensional structure data file (PDB) based on the protein name and performed filtering and clarification operations. After final filtering, we obtained approximately 120k protein sequences and their corresponding three-dimensional structure files.

#### (b) US-align method

The US-align (Universal Structure Alignment) method is a calculation method based on structural alignment, mainly used to compare the similarity between protein three-dimensional structures (*Zhang et al., 2022*). The US-align method can handle protein structures of different lengths and shapes, and has good robustness. The algorithm adopts an iterative optimization strategy, which can converge quickly and is more efficient than the original TM-align method. The TM-align method is also an algorithm based on protein structure alignment, using TM-score as the objective function to evaluate the similarity of protein structures (*Zhang and Skolnick, 2005*). US-align has been improved on the basis of TM-align, expanding its application scenarios while also improving performance.

In our research, we chose US-align as the tool for generating the training dataset. Firstly, this method directly calculates the similarity score TM-score based on the three-dimensional structure, which is currently a relatively new computing tool with high accuracy. Secondly, this tool has good running speed and can help us generate the dataset quickly. The specific flowchart for generating the dataset is shown in Fig. 4.

**Fig. 4.**
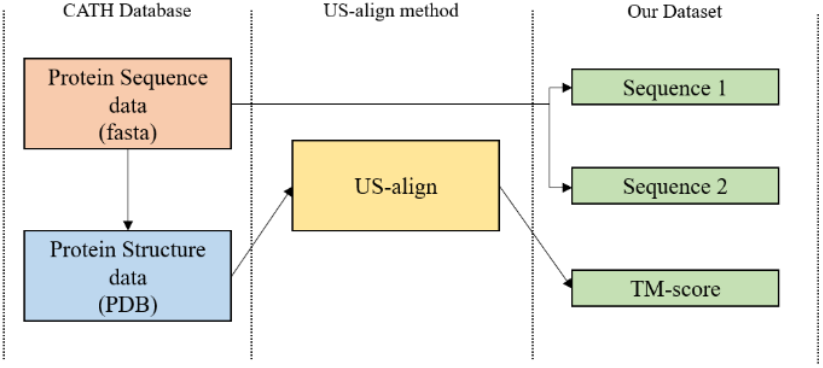
Flowchart for generating datasets

#### (c) Our Dataset

Using the above method, we spent about 3 months generating approximately 1.08B unprocessed datasets with a total file size of approximately 160GB. However, by observing the distribution of the dataset, we can see that the majority of proteins are structurally dissimilar. In order to ensure balance during the training process and avoid training the model on imbalanced datasets, we also need to filter this part of the dataset to make it relatively evenly distributed in the TM-score ranging from 0 to 1. Finally, we obtained the filtered dataset.

In order to facilitate future research in meeting the training needs of different tasks, we have generated three filtered datasets of different sizes, which we refer to CATH_TM_score_S, M and L. The number of data entries and dataset size are shown in Table 1.

**Table 1.** The number of data entries and dataset size.

| Name | S | M | L |
| --- | --- | --- | --- |
| Data Entries | 3.0 M | 8.5 M | 23.3 M |
| Dataset Size | 492 MB | 1.3 GB | 3.6 GB |

During the training process, after multiple adjustments and experiments, we finally decided to divide the ratio of the training dataset to the test dataset into 95%: 5%. The reason for the small proportion of the test set is that our data volume is relatively large. Although the proportion is small, its quantity is sufficient to evaluate the training results of the model.

### 3.2 Result

To evaluate the performance of the model, we will conduct various experiments on our model and comparative models. The devices used are all the same, with a graphics card model of RTX 4070ti 12G, a CPU model of Intel i7 13700k, and a memory size of 32GB. The dataset used is the same test dataset that the model has never learned before. In terms of model comparison, we will compare the publicly available training model of TM-vec, the local dataset training model of TM-vec, and our own model. Since the TM-vec team has not yet released their training dataset, we also need to conduct a local training to ensure fairness. In the following representation, we will use TM-vec-remote to represent the published training model of TM-vec, and TM-vec-local to represent the local dataset training model of TM-vec.

#### (a) Results of all range of TM-score intervals

In order to enable the model to encode protein sequences more accurately and have good recognition ability for protein sequences. We hope that the model can have good performance in all TM-score intervals.

If the model can only achieve good prediction results in a small part of the intervals and cannot predict well in the vast majority of intervals, then we do not believe that the model has learned the features of protein sequences well, but only learned the features in the local range. Therefore, in the final result validation stage, we will first evaluate the model in all intervals.

According to the data in Table 2, we can see that our model has only 41% of the parameter count of TM-vec, making it a more lightweight model than TM-vec. In addition, our model has an average error of 0.0561 across all TM-score intervals, which is better than the performance of the comparison model TM-vec, whether it is TM-vec-remote or TM-vec-local. Fig. 5 shows the distribution of error within the specified interval. If the model performs better, we hope that the first item, which has an overall error of less than 0.05, will have a higher proportion of data points, while at the same time, the fewer points with an error greater than 0.05, the better. We can see that our model has a higher proportion in the interval with small errors, which once again proves the superiority of our model.

**Table 2.**
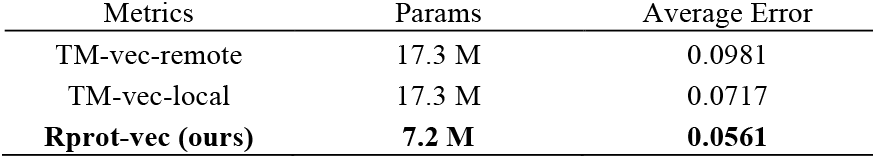
Performance comparison within the all TM-score range.

**Fig. 5.**
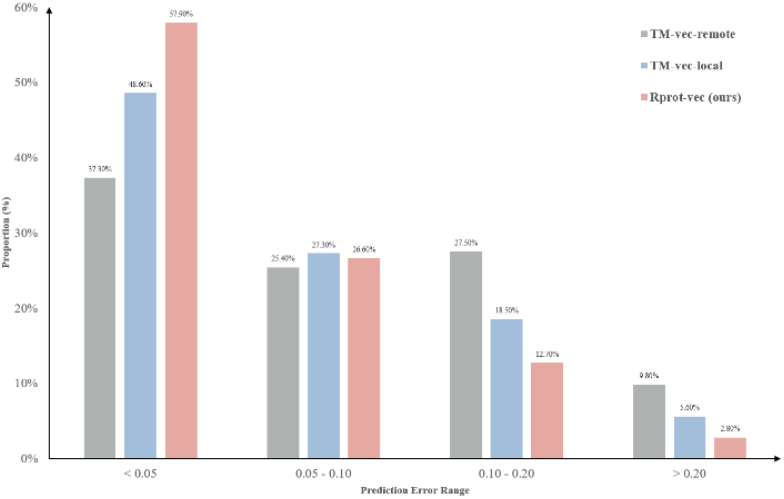
Error performance of different models in all TM-score intervals

In addition, the TM-vec models in Fig. 6 and Fig. 7, as well as our model in Fig. 8, use scatter plots to plot each test data point on the coordinate axis, with different colored points representing different error intervals, which can display the results more clearly.

**Fig. 6.**
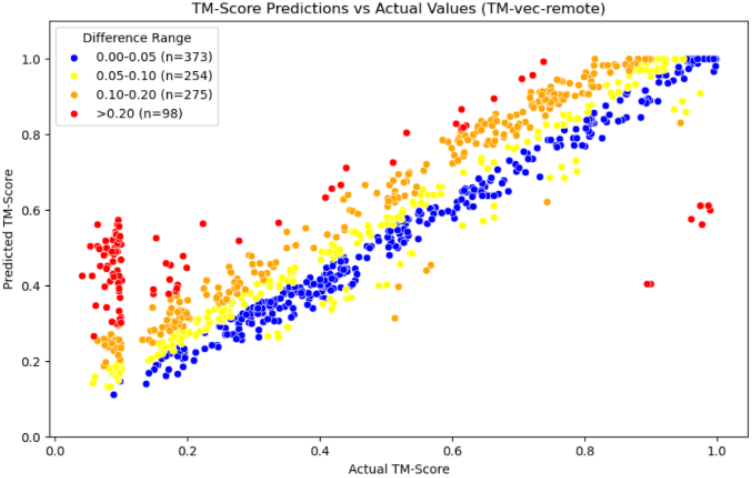
Scatter plot of TM-vec-remote

**Fig. 7.**
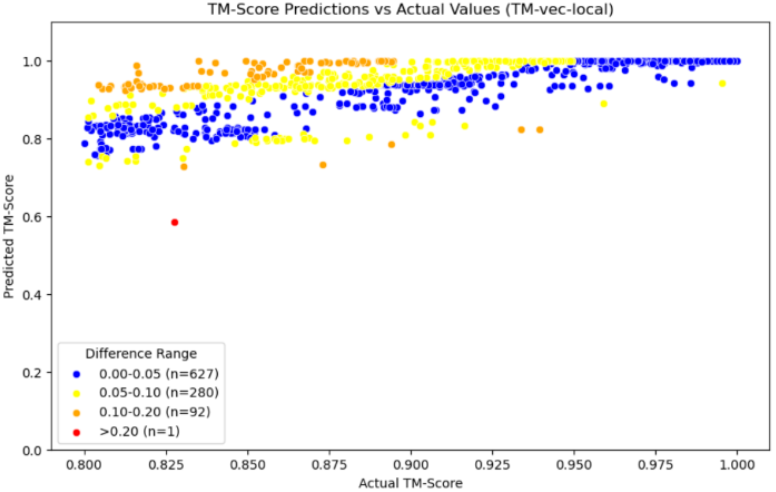
Scatter plot of TM-vec-local

**Fig. 8.**
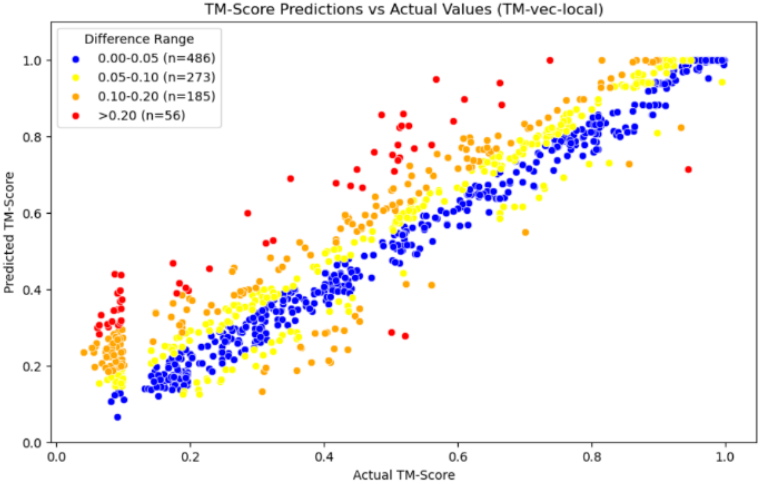
Scatter plot of Our Model

From the comparison of the three graphs above, we can see that the TM-vec model has poor predictive ability when two proteins are completely dissimilar. It is easy to predict two completely dissimilar proteins as partially similar results, while our model has good performance in this interval. In addition, our model also has good prediction results in other intervals.

#### (b) Evaluation of homologous protein detection results

In the introduction section, we mentioned that when TM-score is greater than 0.8, the posterior probability of two proteins in the same folding family is close to 1.0, which means that the two proteins can almost be regarded as homologous proteins. We call this region the homologous region, and the performance of the model in the homologous region will be a very important part, which is related to the ability of the model to identify homologous proteins and whether it can recognize homologous proteins. Therefore, we need to re-evaluate the results of the model on the homologous region.

According to the data in Table 3, we can see that the average error of our model within the homologous region is 0.0438. Better than the performance of the comparison model TM-vec, whether it is TM-vec remote or TM-vec local. In addition, from Fig. 9 we can see that the accuracy prediction rate of our model can reach 65.3%, and there is more data that can be accurately predicted. If TM-vec remote is used, there will be more outliers, with a failure prediction rate (Error > 0.2) as high as 5.7%, and more homologous proteins will not be recognized. We also plotted scatter plots for the homologous region, and Figures 10‒12 are the scatter plot of TM-vec-remote, TM-vec-local and our model in the homologous region.

**Table 3.** Performance comparison within the homologous region.

| Metrics | Params | Average Error |
| --- | --- | --- |
| TM-vec-remote | 17.3 M | 0.0738 |
| TM-vec-local | 17.3 M | 0.0462 |
| **Rprot-vec (ours)** | **7.2 M** | **0.0438** |

**Fig. 9.**
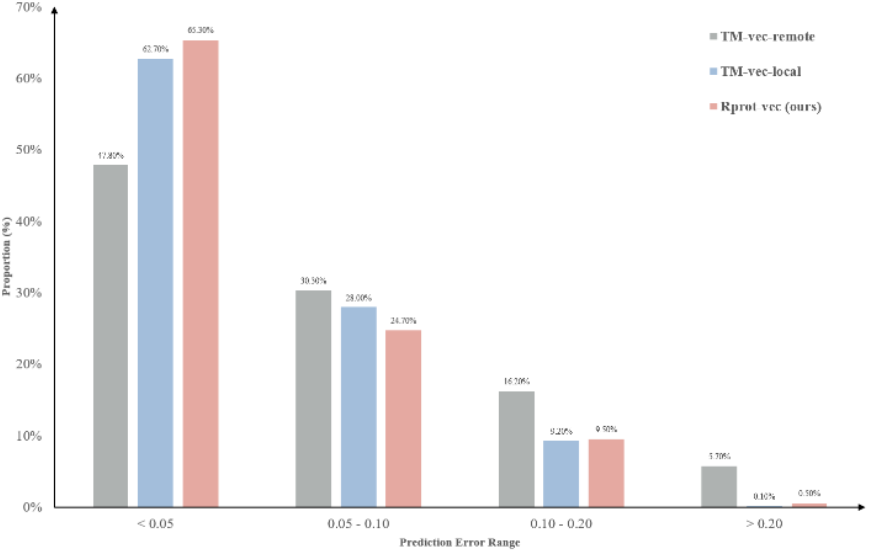
Error performance of different models in homologous region

**Fig. 10.**
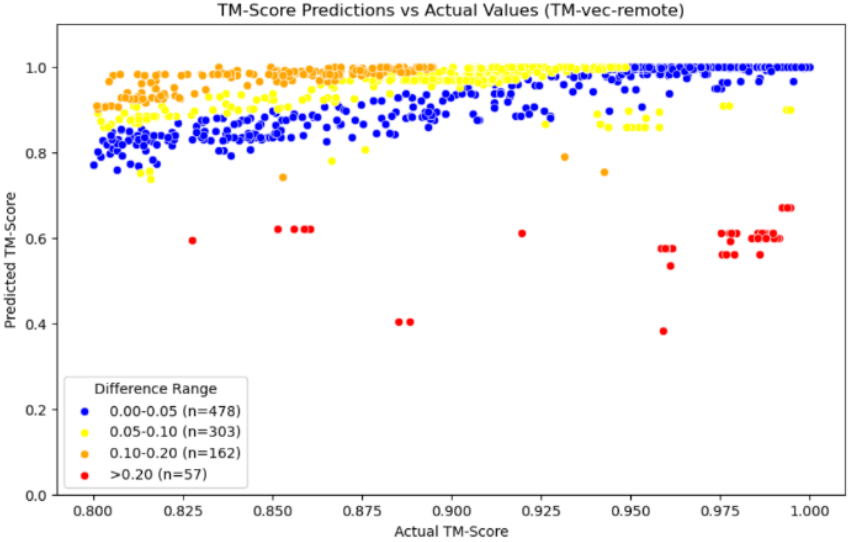
Scatter plot of TM-vec-remote

**Fig. 11.**
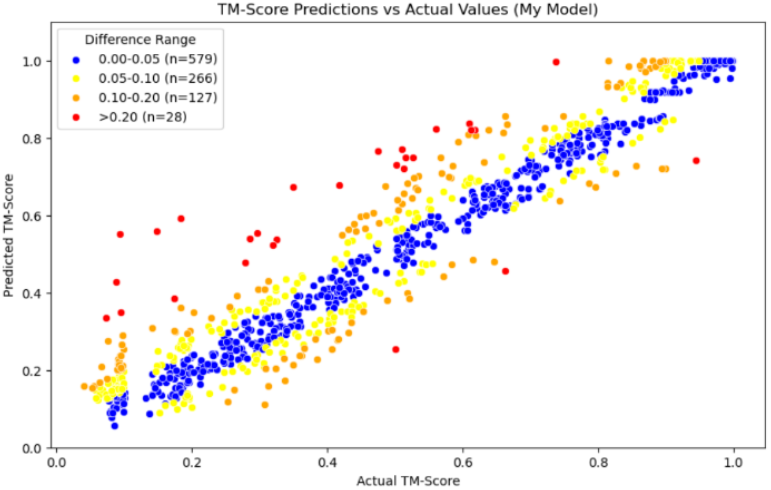
Scatter plot of TM-vec-local

**Fig. 12.**
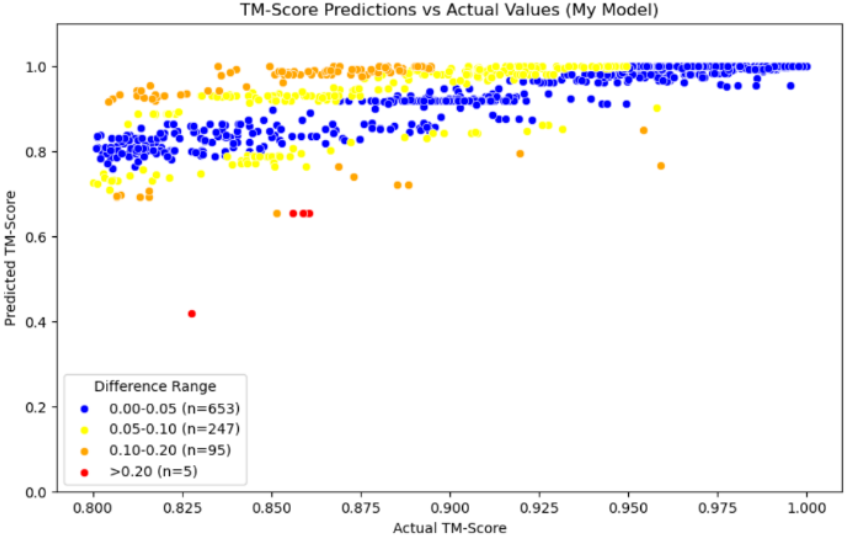
Scatter plot of Our Model

### 3.3 Application

We know that when two proteins have similar three-dimensional structures, their functions are likely to be similar. In terms of application, our model can be utilized in many studies that use structure to infer function. Here, I will list the most typical application case, which is the research on protein drug repositioning and the discovery of novel drugs.

By efficiently calculating the similarity between protein drug sequences, the model can quickly identify the homologous relationship between protein drug pairs. If one protein drug A and another protein drug B have highly similar structures, we can make a preliminary inference that A and B have similar efficacy. The use of this model can enable existing drugs to be used for new therapeutic targets, significantly accelerating the research process of drug repositioning.

In the simulation experiment, we used partial protein drug sequences from the DrugBank database (*Wishart et al., 2018*) (*Knox et al., 2024*) to compare the structural similarity of these drugs pairwise. Finally, we plotted a heatmap of the comparison results, as shown in Fig. 13, where the redder the structure, the more similar it is, and the bluer the structure, the less similar it is.

**Fig. 13.**
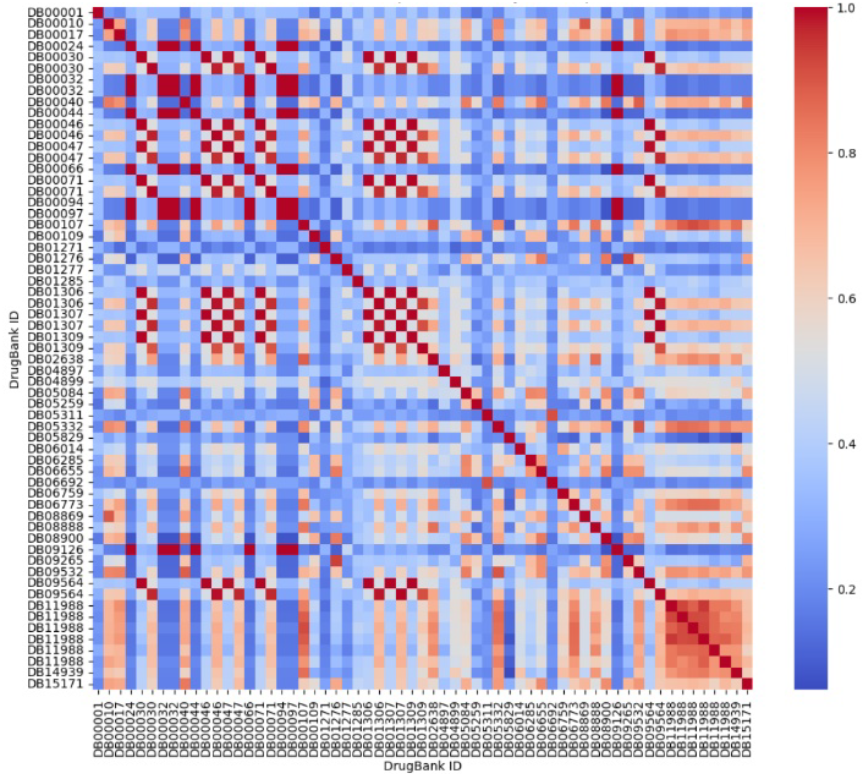
Heatmap of protein drug comparison

In this figure, we have found an interesting pair of drugs, which are DB05311 (Ecallantide) and DB06692 (Aprotinin), In the figure, we can see that these two drugs are not only identical to themselves, but also highly structurally similar to each other. In addition, they exhibit structural dissimilarities with the remaining protein drugs. At this point, we can open the DrugBank database to study this pair of drugs.

The summary of the protein drug Ecallantide is as follows: “Ecallantide is a kallikrein inhibitor used to prevent and treat acute attacks caused by Hereditary Angioedema (HAE).” (DrugBank, 2024) (*Knox et al., 2024*). The summary of Aprotinin is: “Aprotinin is a serine protease inhibitor used to reduce the risk for perioperative blood loss and the need for blood transfusion in high-risk patients during cardiopulmonary bypass for coronary artery bypass graft surgery.” (DrugBank, 2024) (*Knox et al., 2024*).

We found that both exert their effects by inhibiting the activity of serine proteases, but Ecallantide is more targeted and has strong specificity, while Aprotinin has broad spectrum and may have potential in a wider range of diseases, but may also bring some side effects.

By using our model, we can quickly calculate and screen proteins with similar structures. After screening, we can further study the repositioning of protein drugs through in vitro experiments, disease models, or clinical trials.

In addition, it is not only possible to achieve this by comparing the structural similarities between known drugs, but also by comparing the protein drug sequences of other proteins with known functions, in order to explore potential functions. This method can efficiently screen candidate molecules with therapeutic potential, thereby accelerating the development process of new drugs. As shown in Fig. 14, we randomly selected 100 proteins from the CATH database and compared them with some of the selected protein drugs to draw a heatmap. The redder the dots, the more noteworthy the parts, indicating that there is a high probability of structural similarity.

**Fig. 14.**
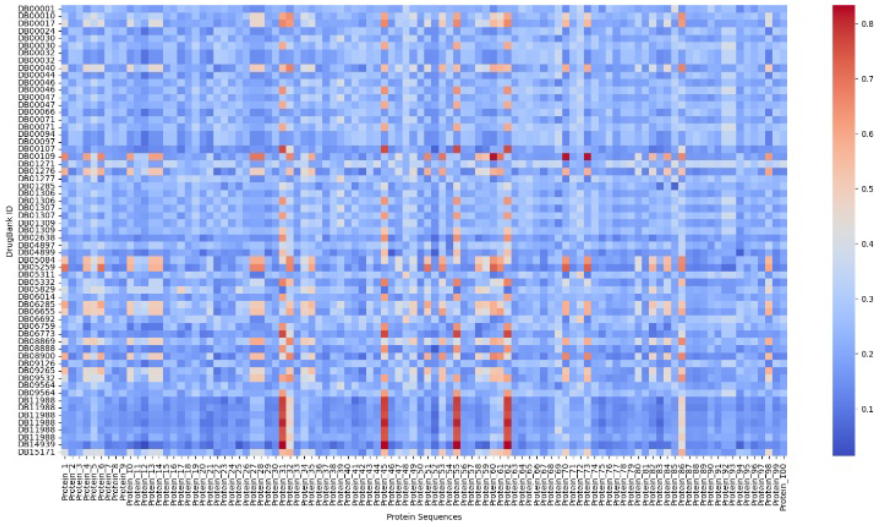
Heatmap of protein drugs and other proteins

## 4 Conclusion

In our study, we proposed an improved method Rprot-vec that can calculate structural similarity using only protein sequences, without the need to calculate through three-dimensional structures. Rprot-vec combines global and local features to fully extract key feature information from protein sequences, models protein sequences and quickly calculates the score of structural similarity.

The experimental results show that Rprot-vec can achieve good results even when trained on small datasets. We trained the model using the CATH_TM_score_M dataset with a size of 1.3GB. Rprot-vec showed better results in both the all TM-score interval and the homologous region. Under the same dataset and training environment, Rprot-vec outperformed the comparison model by 21.8% and 5.19% in both the all TM-score interval and the homologous region.

The structure of Rprot-vec adopts the ProtT5 model as the upstream pre-training encoding model, which first encodes the protein sequence and extracts preliminary feature information. Then, the bidirectional GRU combined with the Attention method is used to further extract global features and attempt to capture key information in the sequence. Next, multiscale convolution is used to capture local features in the protein sequence, and finally, the model calculation of the protein sequence is completed through the output module.

By using Rprot-vec, it is possible to quickly and accurately calculate the structural similarity between two proteins. We look forward to the application of Rprot-vec in the search for homologous proteins. By using Rprot-vec to pre-code a large amount of protein data with known sequences, we can construct the required protein coding database. In subsequent searches, we only need to encode the search sequence and calculate the cosine similarity between the encoded search sequence and the searched sequence in the coding database one by one. The speed of calculating cosine similarity is much faster than that of most deep learning models. This method can help us quickly find proteins in the database that have similar structures to the search sequence.

## Notes

### Competing Interest Statement

The authors have declared no competing interest.

